# An IL-12 mRNA-LNP adjuvant enhances mRNA vaccine induced CD8^+^ T cell responses

**DOI:** 10.1101/2024.07.29.605626

**Authors:** Emily A. Aunins, Anthony T. Phan, Mohamad-Gabriel Alameh, Elisa Cruz-Morales, David A. Christian, Molly E. Bunkofske, Garima Dwivedi, Ross Kedl, Drew Weissman, Christopher A. Hunter

**Author notes:** These authors contributed equally (Emily A. Aunins, Anthony T. Phan).

## Abstract

The design of vaccines that induce CD8^+^ T cell responses has historically been a challenge, but the development of viral vectors or lipid nanoparticles (LNPs) to deliver mRNA that encode for target antigens provide more effective strategies to generate protective CD8^+^ T cell memory. Because interleukin-12 (IL-12) supports CD8^+^ T cell expansion and acquisition of effector functions, studies were performed to assess its contribution to the ability of an mRNA vaccine to promote CD8^+^ T cell responses. *In vitro* and *in vivo*, mRNA-LNPs did not stimulate myeloid cell production of IL-12, and the CD8^+^ T cell response to vaccination with the model antigen ovalbumin (OVA) was IL-12 independent. However, co-administration of IL-12 mRNA-LNPs with OVA mRNA-LNPs enhanced OVA-specific CD8^+^ T cell expansion, improved acquisition of effector function and resulted in an expanded memory CD8^+^ T cell pool. These heightened responses were associated with improved protective responses against *Listeria monocytogenes-OVA* and B16 FO-OVA melanoma. Thus, modification of mRNA vaccine formulations by inclusion of a cytokine mRNA provides a strategy to enhance CD8^+^ T cell mediated protection.

## Introduction

Lipid nanoparticles (LNPs) that encapsulate mRNA that encode target antigens provide a modular platform which formed the basis for the development of several vaccines against SARS-CoV-2 that generate robust humoral immunity and anti-viral CD8^+^ T cell responses.^1-5^ However, mRNA vaccine induced humoral immunity against SARS-CoV-2 wanes, and outgrowth of viral variants in response to selective pressure by antibody has led to widespread breakthrough infection in the vaccinated population.^5-7^ The observation that vaccinated individuals remain protected from severe disease despite reduced antibody titers indicates that other layers of immunity, like CD8^+^ T cells contribute to viral control.^8-10^ While the ability to generate CD8^+^ T cell responses to vaccination has previously been a challenge,^11-13^ mRNA-vaccine induced CD8^+^ T cell responses are associated with protection against severe COVID-19 infection,^4,5^ and these memory CD8^+^ T cells are poised to provide long-lasting and cross-variant protective immunity.^2,4,14-16^ Nevertheless, while current mRNA-LNP vaccines result in expansion of spike specific CD4^+^ and CD8^+^ T cells that persist at detectable levels for at least 6 months, they also eventually wane.^17^ In mice, the ability of mRNA-LNP to induce the production of type I interferon is important for the expansion of COVID-specific CD8^+^ T cells.^14^ Little else is known about how the adjuvanticity of mRNA-LNP vaccines contributes to CD8^+^ T cell responses or how this approach can be engineered to generate a larger and more diverse CD8^+^ T cell memory pool.

The acute response to infection typically generates CD8^+^ T cell subsets that include highly differentiated effectors and less differentiated memory-precursor populations, and following resolution of infection, a memory T cell population remains.^18-20^ In contrast, in a murine model of mRNA vaccination, the initial expansion of vaccine-specific CD8^+^T cells is characterized by a memory precursor like phenotype,^2,8^ associated with a long-lived memory CD8^+^ T cell pool.^21^ In several models of vaccination or infection, the size of the initial CD8^+^ T cell response positively correlates with the magnitude of the memory pool following contraction.^22^ Likewise, the ability to protect against challenge is a function of the size of the memory CD8^+^ T cell pool, as well as the phenotype and location of these CD8^+^ T cells.^23-29^ It seems likely that the ability to modify mRNA vaccination approaches could prove useful to generate a larger more heterogeneous effector response that in turn may give rise to a long lived, more efficacious poly-functional memory pool.

IL-12 is a hetero-dimeric cytokine composed of IL-12p40 and IL-12p35 subunits that is produced by monocytes, macrophages, and dendritic cells in response to pattern recognition receptor (PRR) signaling.^30,31^ IL-12 signals through the IL-12 receptor, which is expressed by natural killer (NK) cells and T cells, and during certain infections the production of IL-12 induces these cells to proliferate and produce IFN-γ.^32-34^ Given these properties, IL-12 has been utilized as a vaccine adjuvant to promote cell-mediated immunity against a variety of parasitic, bacterial and viral infections as well as cancer.^35-39^ However, some studies have shown that the ability of IL-12 to drive CD8^+^ T cells toward terminal effector differentiation is detrimental to the development of memory CD8^+^ T cells.^40-42^ Whether IL-12 is produced in response to mRNA-LNP vaccination or influences the subsequent immune response is unclear. Here, studies were performed to assess the role of endogenous IL-12 in the ability of mRNA vaccines to generate a CD8^+^ T cell response and to determine if the inclusion of mRNA that encodes for IL-12 can be used to modulate the magnitude and effector functions of these CD8^+^ T cells.

## Results

### Role of endogenous IL-12 in CD8^+^ T cell responses to vaccination

Macrophages are a potent source of IL-12, and in response to mRNA vaccination these cells efficiently take up mRNA-LNPs and become activated in draining lymph nodes.^14,43^ To assess whether mRNA-LNPs induce IL-12, bone-marrow derived macrophages (BMDMs) were incubated with empty LNPs or mRNA-LNPs that contain either unmodified mRNA (a potent activator of intracellular sensors) or Nl-methyl-pseudouridylated (Ψ) mRNA overnight and assessed for production of IL-12p40 and IFN-α. As expected, unmodified mRNA-LNPs induced high levels of IFN-α, while Ψ-mRNA-LNPs led to modest induction of IFN-α, but neither of these stimuli resulted in robust IL-12 production (Fig. 1A). Likewise, when mice were injected intramuscularly (i.m) with mRNA-LNP, analysis of these cytokines in the draining popliteal and inguinal lymph nodes highlighted that Ψ-mRNA-LNP resulted in induction of IFN-α, but IL-12p40 was unchanged from basal levels.

**Figure 1.**
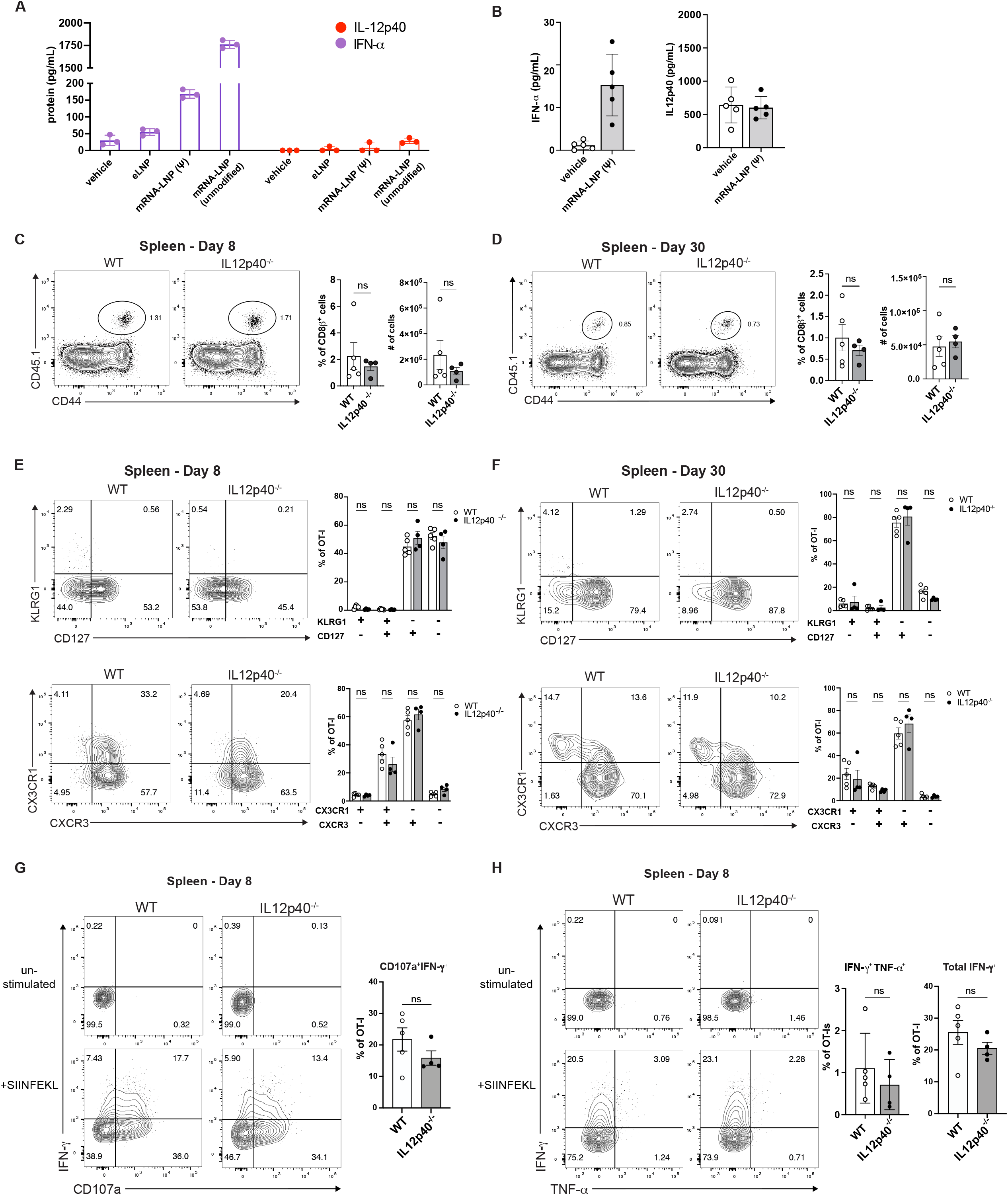
IL-12 is not induced by mRNA-LNPs, and it is not required for the CD8^+^ T cell response to mRNA vaccination. **(A)** C57BL/6 bone marrow derived macrophages (1.5×10^5^ cells/well) were incubated with empty LNP (eLNP), LNP containing pseudouridylated mRNA (Ψ) or LNP containing unmodified mRNA (unmodified) for 20 hours. Cell lysates and their supernatants were assayed for Type I IFN (purple) and IL12p40 (red) protein production. **(B)** C57BL/6 mice were immunized i.m. with mRNA-LNP (Ψ). 6 hours post immunization (hpi), cell lysates generated from draining inguinal and popliteal lymph nodes assayed for IL-12 and IFN -α protein. **(C-F)** Congenically distinct (CD45.1.2) OT-I T cells were adoptively transferred into WT (CD45.2) and IL12p40^-/-^ (CD45.2) mice 1 day prior to immunization. Mice were immunized i.m. with 1 Lig of mRNA-LNP (Ψ) encoding for ovalbumin. Representative flow cytometry plots and summary data show expansion of OT-I T cells by frequency and total cell number in WT (open circles) and IL12p40^-/-^ (closed black circles) mice **(C)** 8 and **(D)** 30 days post immunization. Expression of surface markers of splenic donor OT-I T cells that identify differentiating CD8^+^ T cell subsets (KLRG1 vs CD127 and CXCR3 vs CX3CR1) is shown in representative flow cytometry plots and summary data at **(E)** 8 **(F)** 30 days post immunization. **(G-I)** Splenocytes from vaccinated mice were isolated 8 days post immunization and restimulated with SIINFEKL peptide for 4 hours prior to intracellular cytokine staining. Representative flow cytometry plots and summary data exhibit **(G)** IFN-γ production and degranulation (CD107a) staining, **(H)** IFN-γ and TNF-α staining, and **(I)** Frequency of total IFN-γ expressing cells. All summary data plots display the mean and standard error of the mean. A Shapiro-Wilk normality test was carried out prior to testing for (C-H). A Mann-Whitney rank-sum test was used for non-normally distributed data for (C-D and G). A two way-ANOVA with post-hoc Sídák’s multiple comparison test was performed for (E-F). Data shown are from one representative experiment performed 2 times for (A-B), and 3 times for (C-H).

To determine whether endogenous IL-12 contributes to the CD8^+^ T cell response to mRNA vaccination, OT-I CD8^+^(OT-I) T cells that recognize SIINFEKL, the immunodominant CD8^+^ T cell epitope of Ovalbumin (OVA), presented on the H2-K^b^ allele of major histocompatibility complex class I (MHC-I), were transferred into wildtype (WT) or IL12p40 deficient (IL12p40^-/-^) hosts prior to immunization with OVA-encoding mRNA-LNPs (LNP-OVA). At days 8 and 30 post immunization the expansion, phenotype, and function of the OT-I T cells in the draining inguinal and popliteal lymph nodes (dLNs), spleen, and lung were analyzed. In these tissues in WT and IL12p40^-/-^ mice, there was a comparable expansion of OT-I T cells (Fig. 1C&E, Supp. Fig. 1A). Next, a panel of surface receptors (KLRG1, CD127, CXCR3, CX3CR1) was used to assess the differentiation and activation status of the OT-I T cells at early and late time points. IL-12 promotes expression of killer cell lectin type receptor G1 (KLRG1), (a marker of highly differentiated effector CD8^+^ T cells^18,21,34^) and when used in combination with CX3CR1 can be used to identify functionally distinct populations of effector CD8^+^ T cells.^18,44^ Following vaccination of WT animals, a KLRG1^lo^CX3CR1^lo^CXCR3^hi^CD127^int^ phenotype dominates, an indication that these cells are not highly differentiated effectors but rather possess a memory precursor phenotype. In WT or IL12p40^-/-^ hosts there was no difference in the expression of these markers by OT-I T cells at days 8 or 30 post immunization (Fig. 1D&F). Likewise, at day 8 post-immunization, the absence of IL-12 did not impact the ability of OT-I T cells to produce IFN-γ and TNF-α and degranulate (as measured by surface expression of CD107a) (Fig. 1G-H). In similar experiments performed in the absence of OT-I T cell transfer, use of an H2-K^b^ tetramer loaded with SIINFEKL allowed identification of endogenous OVA-specific CD8^+^ T cells, and absence of IL-12 did not impact these responses (supp. Fig. 1B). In other model systems, the ability of CD4^+^ T cells to license type I conventional dendritic cells (cDC1s) to produce IL-12 contributes to CD8^+^ T cell activation (Supp. Fig. 1C&D).^45,46^ However, following LNP-OVA vaccination, the absence of CD4^+^ T cells, or cDC1 did not impact the ability of CD8^+^ T cells to expand (Supp. Fig. 2C-E). Thus, multiple pathways associated with the production of IL-12 and generation of CD8^+^ T cell responses do not appear to be major contributors for the ability of mRNA-LNP vaccination to induce initial expansion, differentiation, or acquisition of effector function by CD8^+^ T cells.

**Figure 2.**
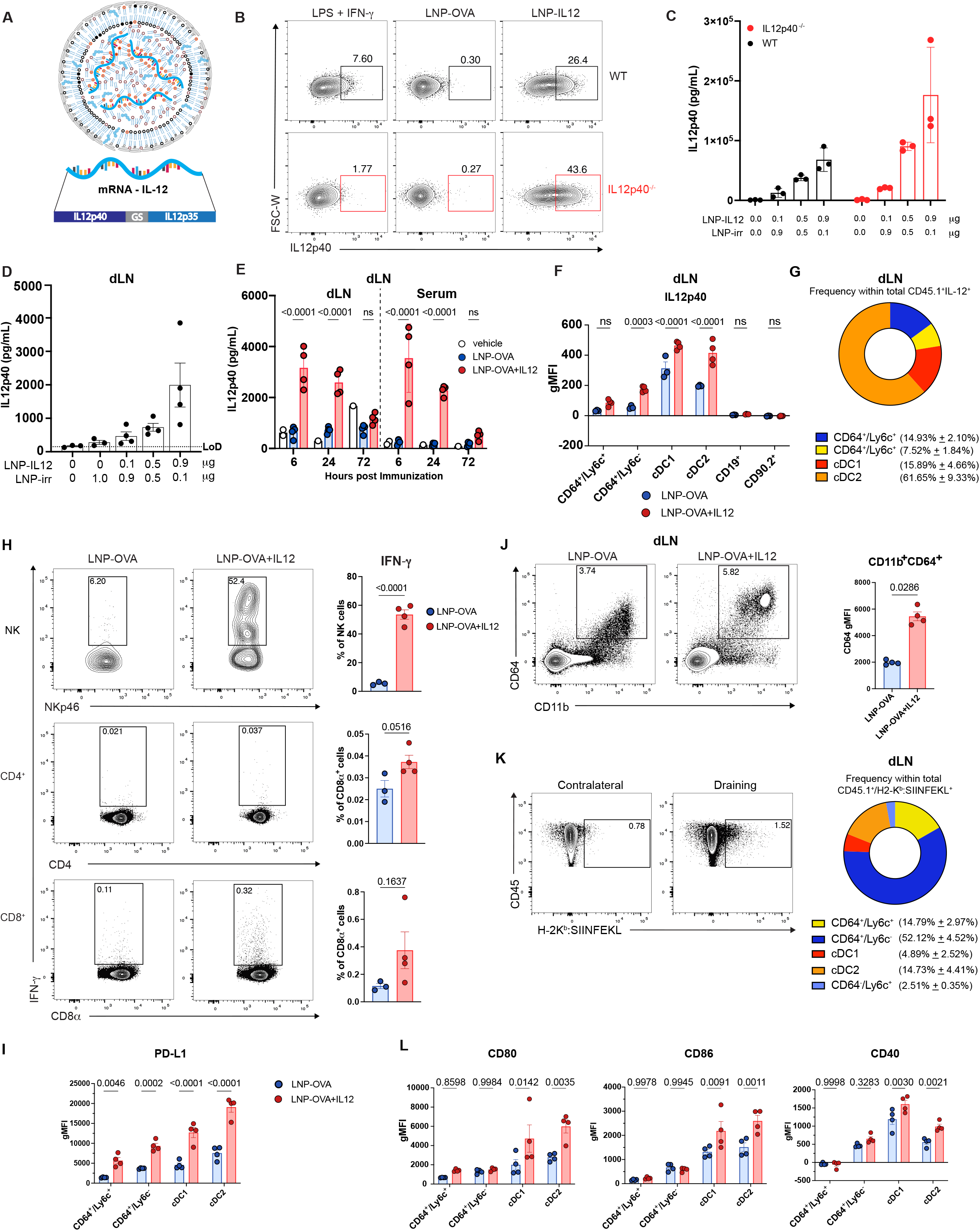
IL-12 mRNA-LNPs induce IL-12p40 protein production. **(A)** Depiction of the LNP-IL12 mRNA construct. **(B)** Flow cytometry analysis of WT (top, black) and IL12p40^-/-^ (bottom, red) BMDMs incubated with LNP-IL12, LNP-OVA, or LPS and IFN-γ for 16 hours prior to addition of protein transport inhibitor. Proportion of IL12p40^+^ BMDMs is indicated in representative flow cytometry plots. **(C)** IL12p40 production detected by ELISA from supernatants of WT and IL12p40^-/-^ BMDMs incubated with increasing doses of LNP-IL12 for 16 hours. **(D)** C57BL/6 mice were immunized i.m. with increasing doses of LNP-IL12, or LNP-irr. 24 hpi, lysates of draining inguinal and popliteal lymph nodes were generated and assayed for IL-12 production by ELISA. **(E)** C57BL/6 mice were immunized i.m. with 1 µg LNP-OVA (blue) or LNP-OVA+IL12 (red). Lysates of draining inguinal and popliteal lymph nodes were generated and assayed for IL-12 production by ELISA at indicated time points. **(F-I)** Flow cytometry analysis of WT mice immunized with LNP-OVA (blue) or LNP-OVA+IL12 (red) and sacrificed 18 hpi. **(F)** IL-12 production (geometric mean fluorescence intensity (gMFI)) by indicated subsets of myeloid lineage cells (CD45.1^+^, CD90.2^-^/B220^-^)(**G**) Parts of whole analysis of IL12p40^+^ cells from mice immunized with LNP-OVA+IL12 (cDC1: CD45.1+/B220-→ CD5^-^ → CD64^-^ → Ly6c^-^/Ly6g^-^ → MHC-II^hi^/CD11c^+^ → XCR1^+^/CD172^-^, cDC2: CD45.1^+^/B220’ → CD5^-^ → CD64^-^ Ly6c^-^/Ly6g^-^ → MHC-II^hi^/CD11c^+^ → XCR^-^/CD172^+^) **(H)** Representative flow cytometry plots and summary data IFN-γ production by NK cells (CD90.27B220^-^ → NK1.17NKp46^+^), CD8^+^ T cells (CD90.27B220^-^ → NK1.17^+^/NKp46^+^ → CD87CD4^-^), and CD4^+^ T cells (CD90.2^+^/B220’ → NK1.1^+^/NKp46^-^ → CD87CD4^+^) **(I)** Summary data of PD-L1 expression by gMFI on myeloid cell populations. **(J-L)** Flow cytometry analysis of myeloid subsets, antigen-presentation, and costimulatory molecular expression of WT mice 72 hpi with LNP-OVA (blue) or LNP-OVA+IL12 (red). **(J)** Analysis of CD64 and CD11b expression by myeloid cells. **(K)** H2-K^b^:SIINFEKL antigen presentation staining on indicated myeloid cells as subsetted above. **(L)** Expression of costimulatory and activation markers of indicated myeloid subsets. All summary data plots display the mean and standard error of the mean. A Shapiro-Wilk normality test was carried out prior to testing for (H, J) followed by a standard unpaired student’s T-test. A two way-ANOVA with post-hoc Sídák’s multiple comparison test was performed for (E-F, I, & L). Data shown are from one representative experiment performed 2 times for (B-C), 1 time for (D&G) and 3 times for (F-L).

### IL-12 mRNA-LNPs expand the CD8^+^ T cell response to mRNA vaccination

Although endogenous IL-12 is not required for the CD8^+^ T cell response to LNP-OVA vaccination, studies were performed to assess whether the inclusion of an IL-12 mRNA-LNP would alter T cell responses. Here, codon-optimized Ψ-mRNA that encodes for the two subunits of IL-12, p40 and p35 was synthesized and encapsulated in LNPs (Fig. 2A). These coding sequences are linked by a flexible glycine-serine (GS) linker, such that the two subunits are co translated and efficiently heterodimerize into functional IL-12.^47^ To test the efficiency of this formulation, WT and IL12p40^-/-^ BMDMs were incubated with LNP-IL12 particles and intracellular IL-12 staining revealed the induction of IL-12 production and the dose-dependent secretion of IL12p40 (Fig. 2B&C). When WT mice were immunized i.m. with LNP-IL12, a dose dependent translation of IL-12 in the dLNs was observed, and dLNs and serum levels peaked at approximately 3.5-4 ng/mL at 6 hours post-treatment which returned to basal levels by 72 hours (Fig. 2D&E). To determine which cell types produce IL-12, IL12p40^-/-^ mice were immunized with 5 µg LNP-OVA+IL12 or LNP-OVA, and 16 hours later cells from the dLNs were incubated with protein transport inhibitors (Brefeldin A and monensin) for 4 hours before staining for surface markers and intracellular cytokine. In response to LNP-OVA+IL12, B cells (CD19^+^) and lymphocytes (CD90.2^+^) did not express IL-12, but IL-12 expression was readily detected in CD64^+^ monocytes (Ly6c^+^) and macrophages (Ly6c) and cDCs (Fig. 2G).

Because vaccination with mRNA-LNP induces the activation and recruitment of immune cells to the dLN,^14,48^ the impact of IL-12 mRNA on these populations was assessed. Most noticeably, while mice immunized with LNP-OVA alone did not produce significant IFN-γ, when these particles were used in combination with LNP-IL12 approximately 50% of NK cells, 0.4% of CD8^+^ and 0.04% CD4^+^ T cells stained positive for IFN-γ (Fig. 2H). At 72 hours post immunization, LNP-OVA alone and LNP-OVA+IL12 were associated with an accumulation of CD64^hi^ macrophages and monocytes in dLNs. While incorporation of LNP-IL12 did not alter the size of the cDC population or the proportion of cDC1 and cDC2 subsets in the dLN (data not shown) there was a marked increase in myeloid cell expression of PD-L1 and CD64 (F_c_γ receptor I), both canonical targets of IFN-γ signaling (Fig. 2I-J). Use of an antibody that recognizes SIINFEKL complexed with the H2-K^b^ allele of MHC class I^49^ revealed that at 72 hours post vaccination the highest proportion of cells presenting SIINFEKL were CD64^+^ macrophages and CD11c^+^ dendritic cells (Fig. 2K). The inclusion of LNP-IL-12 did not impact the numbers or composition of cells presenting antigen but was associated with increased expression of the co-stimulatory proteins CD80, CD86, and CD40 by these APC populations, most notably cDC2s (Fig. 2L). Thus, delivery of LNP-IL12 leads to transient production of IL-12 in dLNs, associated with altered immune cell infiltration, increased production of IFN-γ and activation of local antigen presenting cells.

### IL-12 mRNA-LNPs enhance effector CD8^+^ T cell expansion

To determine if incorporation of LNP-IL12 into mRNA vaccination alters the CD8^+^ T cell response, WT OT-I T cells were transferred into congenically distinct C57BL/6 mice one day prior to vaccination with OVA mRNA-LNPs (LNP-OVA) or LNP-OVA and LNP-IL12 (LNP-OVA+IL12), and the response of OT-I CD8^+^ T cells was tracked in the blood. To ensure equal lipid and antigen dose were given in each group, an empty LNP (eLNP) was mixed with LNP-OVA at the same ratio (1:1 by mass) as LNP-OVA+IL12. Administration of LNP-OVA led to expansion of OT-I CD8^+^ T cells, but in comparison, mice that received LNP-OVA+IL12 had a > 4-fold increase in frequency of OT-I CD8^+^ T cells (Fig. 3A-B). Likewise, the inclusion of LNP-IL12 also led to an increase in the frequency and total number of OT-I T cells in the lungs, draining lymph nodes and spleens at day 8 (Fig. 3C). Similar results were obtained using a tetramer against SIINFEKL to detect endogenous responses to OVA (Supp. Fig. 3A&B). The use *of Uniform Manifold Approximation and Projection* for Dimension Reduction (UMAP) analysis of high parameter flow-cytometry data from concatenated splenic OT-I samples illustrated that at day 8 post immunization, LNP-OVA results in the generation of a homogenous effector OT-I population (KLRG1^lo^CX3CR1^lo^CXCR3^hi^CD127^int^) (Fig. 3D). Overlaying the expression of a number of cell surface markers that define subsets of effector CD8^+^ T cells on the UMAP projection revealed a dichotomy between OT-I T cells from LNP-OVA versus LNP-OVA+IL12 immunized mice (Fig. 3D-F). In mice that receive LNP-OVA+IL12, there was more heterogeneous expression of CXCR3, CX3CR1, KLRG1, CD43 and CD27 in comparison with mice that receive LNP-OVA (Fig. 3D-F). These differences were also reflected in the lung and to a lesser extent, the dLN (data not shown). These expression patterns indicate that the inclusion of IL-12 does not significantly alter the number of OT-I T cells with memory potential (KLRG1^lo^CX3CR1^lo^CXCR3^hi^CD127^int^) but does result in an increased number of more terminally differentiated effector QT-I T cells (CX3CR1^+^KLRG1^+^) (Fig. 3G).

**Figure 3.**
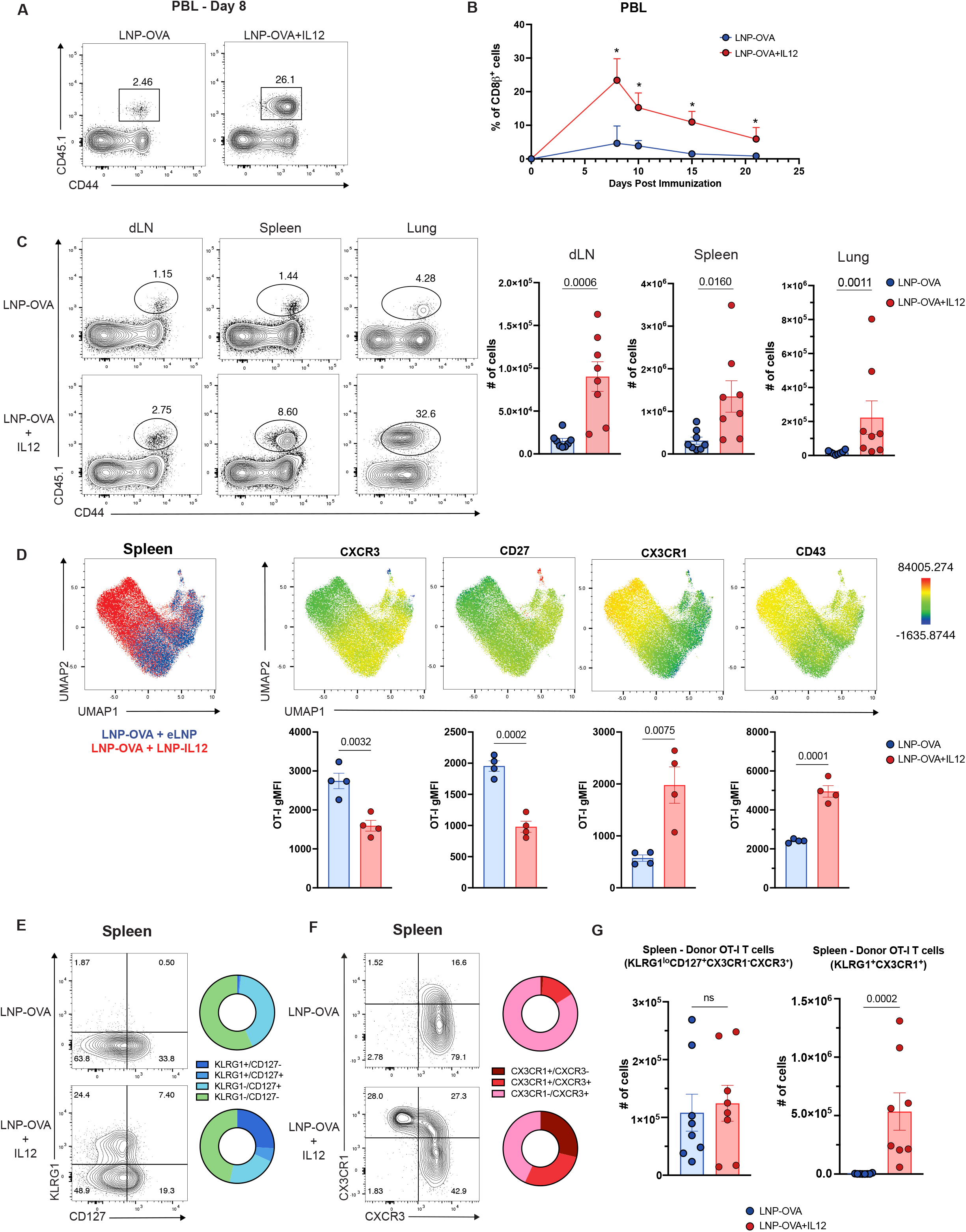
Incorporation of LNP-IL12 enhances vaccine-specific CD8^+^ T cell expansion and alters phenotype. **(A-G)** Flow cytometry analysis of donor OT-I CD8^+^ T cells in indicated tissues of WT (CD45.2) host mice over time that received congenically distinct (CD45.1.2) OT-I CD8^+^ T cells followed by immunization with LNP-OVA (blue) or LNP-OVA+IL12 (red). **(A)** Representative flow cytometry plots of donor OT-I CD8^+^ T cells as a proportion of host CD8^+^ T cells in peripheral blood 8 days post immunization. **(B)** Frequency of OT-I CD8^+^ T cells in blood at days 8, 10, 15, and 21 post immunization. **(C-G)** Flow cytometry analysis of donor OT-I CD8^+^ T cells at day 8 post-immunization in draining LN, Spleen, and Lung. **(C)** Representative flow cytometry plots of the proportion of donor OT-I CD8^+^ T cells and summary graphs of absolute number of OT-I CD8^+^ T cells identified in indicated tissues. **(D)** UMAP projection of high parameter flow cytometry data of donor OT-I CD8^+^ T cells in the spleen 8 days post immunization with LNP-OVA or LNP-OVA+IL12 (top) of phenotypic markers of CD8^+^ T cell effector subsets. Source of donor OT-I CD8^+^T cells overlayed on the projection (top left). Expression pattern of each indicated marker is overlayed on the UMAP projection. Summary gMFI plots of indicated marker by donor OT-I CD8^+^ T cells from their respective host (bottom). **(E-G)** Representative flow cytometry plots with summary parts of whole analysis of donor OT-I CD8^+^ T cells in the spleen. **(E)** KLRG1 vs CD127 **(F)** CX3CR1 vs CXCR3 expression 8 days post immunization. **(G)** Summary plots showing total number of memory precursor like cells (KLRG1 CD127^+^CX3CR1’CXCR3^+^) and effector cells (KLRG1^+^CX3CR1^+^) 8 days post immunization. All summary data plots display the mean and standard error of the mean. A Shapiro-Wilk normality test was carried out prior to testing for (B-D, G) followed by a Mann-whitney rank sum test. Data was pooled from all (2) replicates, except UMAP analysis and summary gMFI which was taken from one representative experiment due to variability in staining intensity.

In some models, the ability of IL-12 to promote CD8^+^ T cell effector responses can be detrimental to the formation of the CD8^+^ T cell memory pool.^40-42^ When the QT-I populations were surveyed > 4 weeks post vaccination these populations had undergone a marked contraction but the inclusion of LNP-IL12 resulted in a larger pool of QT-I CD8^+^ T cells across all tissues examined (Fig. 4A). At memory time points, few QT-I T cells can be found in dLNs for either vaccine group and thus were not analyzed. In contrast to 8 days post immunization, the phenotype of QT-I CD8^+^ T cells from each vaccine group is more similar at these later time points, each group dominated by CD127^+^KLRGT^-^, CXCR3^+^CX3CR1^-^ QT-I T cells (Fig. 4B&C). Nevertheless, in mice that receive LNP-QVA+IL12 there was an expanded CD27^10^ population (Fig. 4D), a phenotype associated with increased killing capacity upon secondary challenge^25^ and a significantly expanded central memory (CD127^+^CD11a^+^CD44^hi^CD62L^+^) and effector memory (CD127^+^CD11a^+^ CD44^hi^CD62L^+^) QT-I CD8^+^ T cell population (Fig. 4E). Moreover, when vaccinated mice were injected intravenously with a fluorescent antibody against CD8α to distinguish vascular and parenchymal populations of CD8^+^ T cells,^50^ LNP-IL-12 was associated with a greater number of parenchymal CD8^+^ T cells (Fig. 4F). Thus, incorporation of LNP-IL12 in mRNA-LNP vaccination enhances the size of the memory CD8^+^ T cell pool across tissues.

**Figure 4.**
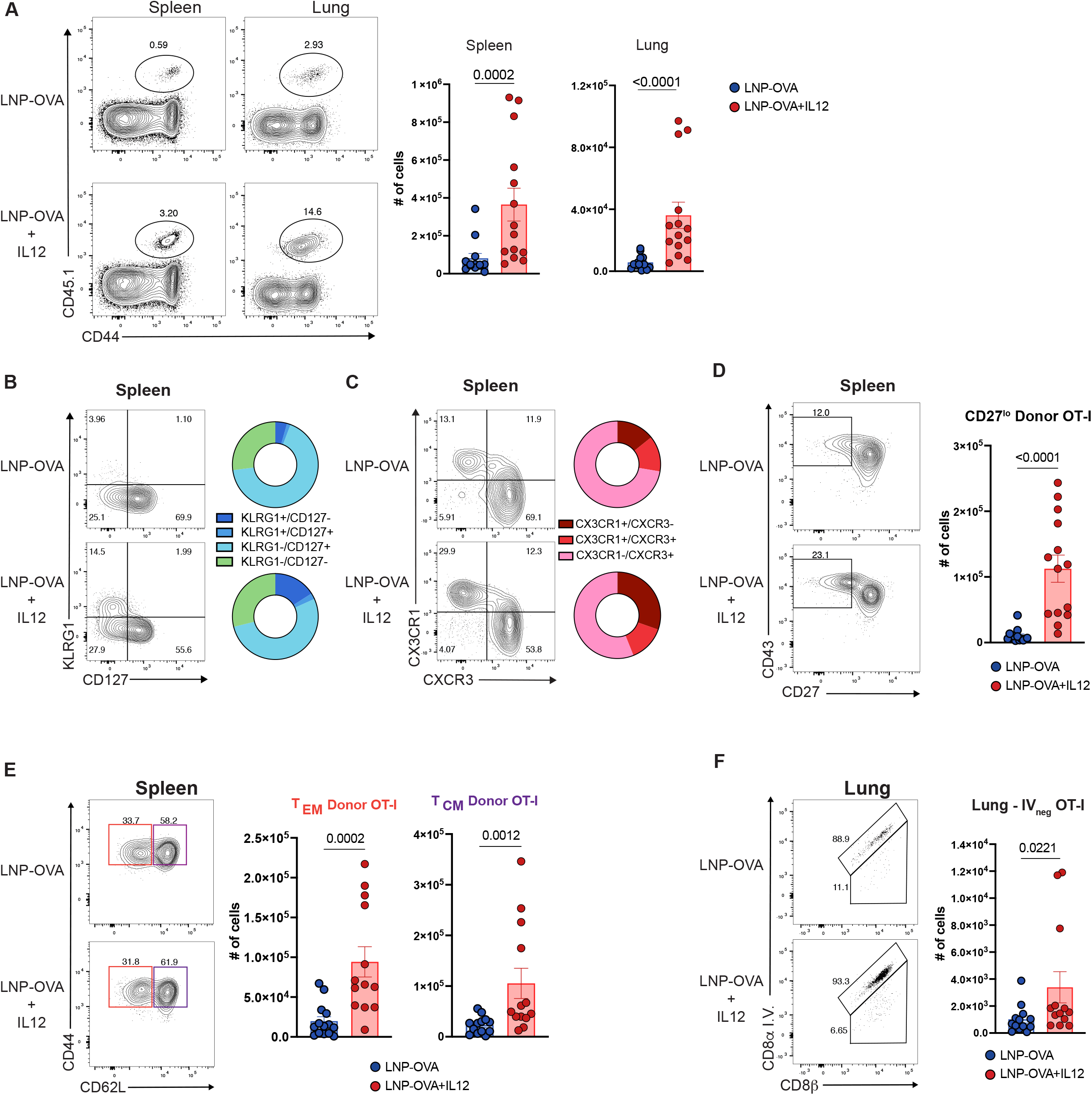
Incorporation of LNP-IL12 into vaccination results in sustained alteration of effector and memory cell differentiation of responding OT-I T cells. **(A-F)** Flow cytometry analysis of donor OT-I CD8^+^ T cells from wildtype host mice immunized with LNP-OVA (blue) or LNP-OVA+IL12 (red) at >28 days post vaccination. **(A)** Representative flow cytometry plots and summary data showing proportion (left) and numbers (right) of donor OT-I CD8^+^ T cell spleen and lung. **(B-C)** Representative flow cytometry plots of phenotypic markers of memory CD8^+^ T cell subsets with summary parts of whole analysis of **(B)** KLRG1 and CD127 and **(C)** CX3CR1 and CXCR3 by donor OT-I CD8^+^ T cells. **(D)** Representative flow cytometry plots and of CD27 and CD43 expression with proportion of CD27^lo^ donor OT-I CD8^+^ T cells indicated and summary graph of total number of CD27^lo^ cells from spleen. **(E)** Central memory (CD127^hi^, CD62L^hi^, CD11a^hi^, CD44^hi^) and effector memory (CD127^hi^, CD62L^lo^, CD11a^hi^, CD44^hi^) QT-I CD8^+^ T cells observed in the spleen. **(F)** Representative flow cytometry plots (left) of IV labeling of CD8^+^ T cells indicating circulating vs. tissue localized QT-I CD8^+^ T cells identified in the lungs post immunization. Summary graph of absolute numbers of IV_neg_ QT-I CD8^+^ T cells observed in the lungs. All summary data plots display the mean and standard error of the mean. A Shapiro-Wilk normality test was carried out prior to testing for (A, D-F) followed by a Mann-whitney rank sum test. Data was pooled from all (3) replicates.

### IL-12 mRNA-LNPs enhance OT-I function

To determine the impact of LNP-IL12 in mRNA vaccination on CD8^+^ T cell function, OT-I CD8^+^ T cells were transferred into mice, and at days 8 and 29 post vaccination, splenocytes from each group were restimulated with SIINFEKL peptide and cytokine secretion and degranulation assessed. At day 8 after immunization, OT-I CD8^+^ T cells from each group produced IFN-γ and degranulated in response to restimulation at a similar frequency, but LNP-IL12 resulted in an increase in the levels of IFN-γ synthesized on a per cell basis (Fig. 5A). Similarly, at 4 weeks post immunization, LNP-IL12 resulted in an increased frequency of OT-I CD8^+^ T cells able to degranulate and produce IFN-γ in response to SIINFEKL (Fig. 5A). Moreover, in an *in vivo* cytotoxic T lymphocyte (CTL) assay in which vaccinated mice were challenged with SIINFEKL pulsed target cells, those that received LNP-OVA killed approximately 30% of the target cells, but the inclusion of IL-12 mRNA doubled the level of target cell killing to approximately 66% (Fig. 5B). Additionally, vaccinated mice were challenged >4 weeks after primary vaccination with an OVA-expressing strain of *L. monocytogenes* (Lm-OVA).^51–53^ Prior to challenge, mice that received LNP-OVA+IL12 had a greater frequency of circulating OVA-specific CD8^+^ T cells (Fig 5C). Following challenge, highest levels of protection were also observed in those mice that received LNP-OVA+IL12, and pre-challenge OVA-specific CD8^+^ T cell frequency in blood correlated with protection (Fig. 5D, Supp. Fig 3A). Furthermore, to determine whether a reduced dose of LNP-OVA+IL12 would still result in enhanced protection in against *L. monocytogenes*, mice were vaccinated with 1 µg LNP-OVA, 1 µg LNP-OVA+IL12, or 0.25 µg LNP-OVA+IL12 and challenged 33 days later with Lm-OVA. Despite more modest differences in circulating OVA-specific CD8^+^ T cells at the time of challenge (Fig. 5E), both doses of LNP-OVA+IL12 were signficiantly more protective than 1 µg of LNP-OVA (Fig. 5F).

**Figure 5.**
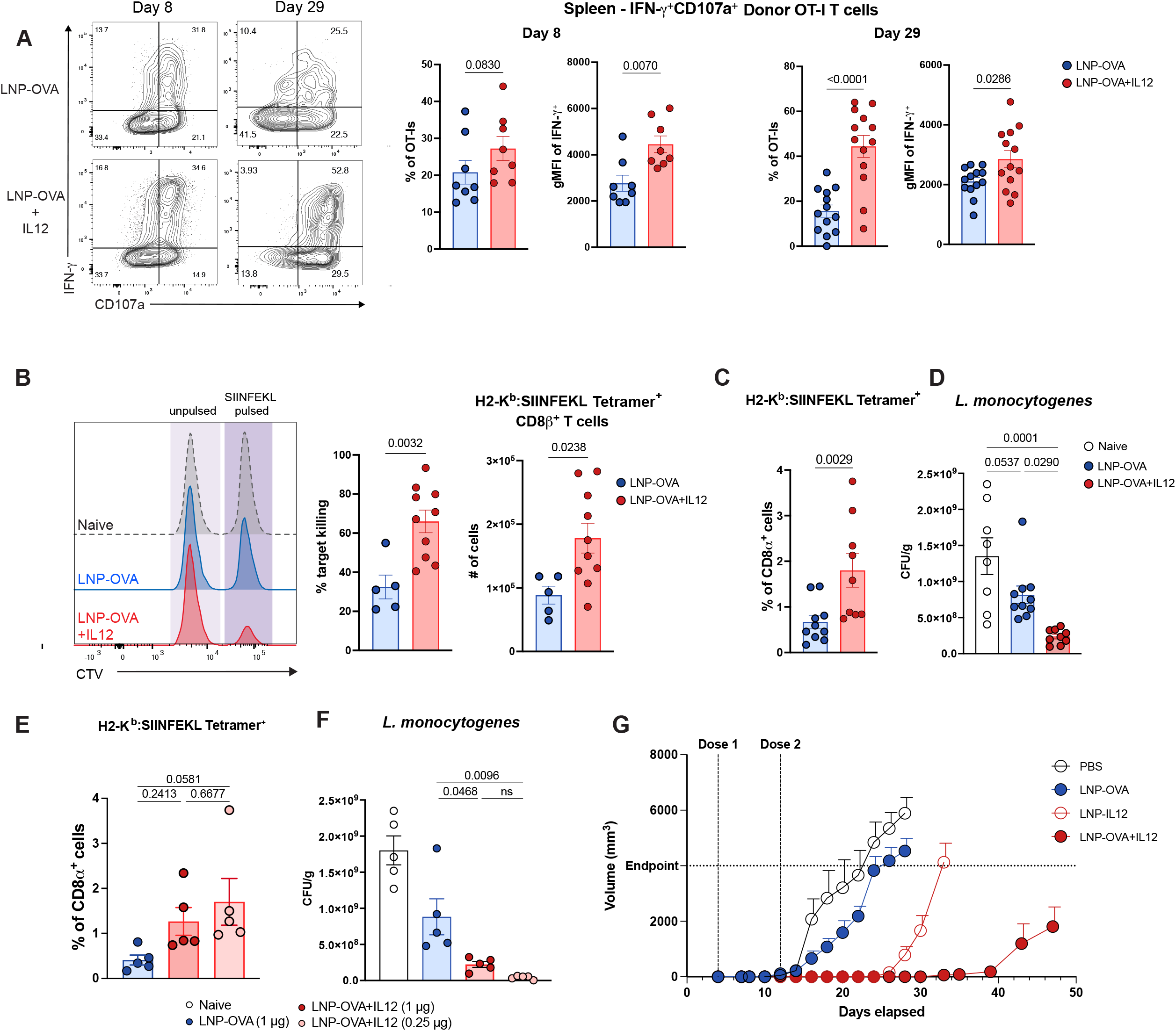
IL-12 enhances effector function and leads to enhanced protection in therapeutic and prophylactic vaccination. **(A)** Flow cytometry of analysis IFN-γ production and degranulation (CD107a) by congenically distinct donor OT-I CD8^+^ T cells in the spleen of wildtype host mice day 8 or day 29-30 post immunization with LNP-OVA (blue) or LNP-OVA+IL12 (red). Representative flow cytometry plots (left), summarized proportion of CD107a^+^ IFN-g^+^ donor OT-I CD8^+^ T cells at indicated time points post immunization (right). IFN-γ expression by IFN-γ^+^ OT-I CD8^+^ T cells also summarized for each time point. **(B)** In vivo cytotoxic lymphocyte assay of WT mice 30 days post immunization with LNP-OVA or LNP-OVA+IL12. Representative flow cytometry plots of target cell killing (left) was analyzed as a function of the ratio of pulsed to unpulsed cells remaining 16 hours post transfer. Summarized target cell killing and number of H2-K^b^ SIINFEKL^+^ CD8^+^ T cells in immunized mice following assay (right). **(C)** Frequency of OVA-specific CD8^+^ T cells in peripheral blood of WT mice immunized with LNP-OVA or LNP-OVA+IL12 >4 weeks post immunization. **(D)** CFU/g of OVA-expressing strain of L. monocytogenes (Lm-OVA) isolated from spleens of WT naive, LNP-OVA immunized, or LNP-OVA+IL12 immunized mice challenged with 2×10^6^ CFU of Lm-OVA 48 hrs prior. **(E)** Frequency of OVA-specific CD8^+^ T cells in peripheral blood of WT mice immunized with LNP-OVA (1 µg)(blue) or LNP-OVA+IL12 (1 µg (dark red circles) and 0.25 µg (light red circles)) >4 weeks post immunization. **(F)** CFU/g of OVA-expressing strain of L. monocytogenes (Lm-OVA) isolated from spleens of WT naive, LNP-OVA immunized (blue), LNP-OVA+IL12 (1 µg) (dark red), LNP-OVA+IL12 (0.25 µg) (light red) immunized mice challenged with 2×10^6^ CFU of Lm-OVA 48 hrs prior. **(G)** Flank tumor growth tracked over time in WT mice that were implanted subcutaneously with 2.5×10^5^ B16F0 melanoma cells engineered to express OVA. At 4- and 12-days post tumor implantation, mice received a 10 µg vaccination (Dose 1 and Dose 2) with the mRNA-LNP indicated (PBS, open circle, LNP-OVA, blue filled, LNP-IL12, open red, LNP-OVA+IL12, red filled). Data from 1 representative experiment, n=10 per group. All summary data plots display the mean and standard error of the mean. A Shapiro-Wilk normality test was carried out prior to testing for (A-C) followed by a Mann-whitney rank sum test. A one-way ANOVA with follow-up Tukey’s test was used for (D-F). Data pooled from all (3) replicates for (A), from all (2) replicates for (C-D), data from 1 replicate performed 1 time (B, E F), from 1 replicate performed 2 times for (G).

Previous studies have shown that mRNA vaccination can be used in a therapeutic setting in a model of B16F0 melanoma in which CD8^+^ T cells can mediate protection.^54,55^ Therefore, to assess the utility of IL-12 mRNA in this system, B16F0 melanoma cells that expressed OVA (B16F0-OVA) were implanted into the flanks of mice. At day 4 post-implantation, mice were vaccinated i.m. in the same side as the tumor with vehicle control, LNP-OVA, LNP-OVA+IL12 or LNP-IL12 alone and a second dose of the same vaccine was given 12 days post implantation.^56^ In mice that received dPBS alone, palpable tumors appeared as early as 8 days post-implantation, and by day 15 all mice in this treatment group had reached the humane endpoint tumor volume (Fig. 5G). In mice that received LNP-OVA, there was a delay in the onset of tumor growth until day 11, but most reached humane endpoint tumor volume by day 20 post implantation (Fig. 5G). In contrast, mice that received LNP-IL12 alone had a marked delay in tumor outgrowth (day 20), but once established, these tumors grew to the humane endpoint size of 4000 mm^3^ within 11 days of onset (Fig. 5G). Only mice that received the combination of LNP-OVA+IL12 had sustained tumor suppression past 30 days post implantation (Fig. 5G).

## Discussion

Here, application of IL-12 mRNA-LNPs to tune CD8 T cell responses demonstrates a strategy in which cytokine mRNA-LNPs can enhance the magnitude and breadth of the immune response to vaccination. The use of Ψ modified mRNA is a critical feature that mitigates the toxicity of mRNA-LNP formulations, but there are adjuvant effects associated with the ionizable lipids used in the LNP formulations to allow cytosolic delivery of mRNA. This combination can engage RIG-I and TLR signaling to mediate the innate production of Type I interferon and IL-6 that contribute to humoral and cellular responses.^1,14,57^ The utility of altering the innate response to mRNA-LNPs was highlighted by recent findings that mRNA-LNPs that encode self-reactive cGAS enhanced vaccine immunogenicity and the resulting cellular immune response.^59^ However, the data presented here indicate that the formulations commonly used for approved mRNA vaccines do not engage innate sensors that lead to the production of IL-12. This result is in agreement with the lack of reports that these mRNA vaccine formulations induce IL-12,^1,14^ and are consistent with the observations that IL-12 appears to be dispensable for ability of mRNA vaccination to generate CD8^+^ T cell responses.

The potent adjuvanticity of pro-inflammatory cytokines like IL-12 has motivated their use to enhance therapeutic immune responses,^59-66^ but their systemic administration in the clinic has resulted in extreme adverse effects.^67^ A number of strategies have been developed to limit these systemic effects by delivering lower doses of cytokines to specific tissues,^68^ and a recent report that engineered IL-12 mRNA transcripts to be restricted to the intramuscular injection site showed that this strategy can amplify the humoral response to the BN162b2 SARS-CoV-2 mRNA vaccine independent of age.^69^ In the present study, the formulation of IL-12 into mRNA-LNPs allowed for transient production of IL-12 by professional antigen presenting cells in draining lymph nodes. Administration of IL-12 mRNA-LNPs at the time of immunization enhanced co-stimulatory protein expression and activation of antigen presenting cells at the site and time of naive T cell priming, which resulted in significantly expanded effector and memory CD8^+^ T cell pools. These CD8^+^ T cell populations mimicked responses observed with live infection and were characterized by a more diverse, protective T cell population that resulted in enhanced CD8^+^ T cell-dependent protection in prophylactic vaccination. Moreover, although IL-12 mRNA induced effector populations were characterized by high KLRG1 expression, which is thought to mark short-lived effector cells that form poor memory,^18,21^ recent studies have highlighted that memory CD8^+^ T cells that express KLRG1 retain plasticity and can contribute to long term protection.^25,27,70^ The utility of the inclusion of IL-12 mRNA-LNPs was further emphasized by the observation that a reduced dose (0.25 µg) of LNP-OVA+IL12 was more protective than 1 µg of LNP-OVA in an *L. monocytogenes* challenge. This finding suggests that this strategy not only allows for bypassing toxicity associated with cytokine delivery but can reduce the amount of mRNA-LNP needed per vaccine dose and thus reduce side effects associated with the LNP vaccination.^71^ Here, the ability to use IL-12 mRNA-LNP as an adjuvant given with an mRNA-LNP that encodes a surrogate tumor antigen highlights the efficacy of this approach.

Though the generation of CD8^+^ T cell response to vaccination has historically been considered a roadblock in the design of some vaccines, there are other challenges that may be overcome by the ability to incorporate other cytokine mRNA-LNPs. For example, the use of cytokines that can influence antibody class switching or CD4^+^ T cell effector phenotype (e.g. IL-4, IL-21) may lead to stronger protection in systems where different forms of immunity are important. In recent years, the field of engineered cytokines has highlighted how their structural modification can be used to bias downstream signaling and biological function. The modularity of the mRNA-LNP platform means that this is a tractable system that would allow expression of these altered ligands and may allow precise tuning of the levels of polarization and survival signals that influence T cell fate. Thus, the efficacy of IL-12 mRNA-LNPs provide an important proof of concept of the opportunity to capitalize on decades of cytokine biology research to design adjuvants from a mechanistically informed perspective and create vaccines and therapeutics that provide potent, long-lasting protection.

## Materials and Methods

### Mice

For all strains of mice used in this study, both male and female mice were used. C57BL/6 mice (6-7 weeks old) were purchased from Jackson laboratories (Bar Harbor, ME, USA) and housed in the University of Pennsylvania Department of Pathobiology vivarium for 1-4 weeks before use. Male and female IL12p40^-/-^ mice were previously purchased from Jackson Laboratories and bred in the University of Pennsylvania Department of Pathobiology vivarium in accordance with institutional guidelines, and they were used between 7-12 weeks of age.

### mRNA design and production

The ovalbumin (OVA) and IL-12 cytokine (IL-12) amino acid sequences underwent codon optimization and GC enrichment using our proprietary algorithm to improve expression and reduce potential immunogenicity of the *in vitro* transcribed mRNA. The codon optimized sequences were gene synthetized by Genscript, cloned into our proprietary *in vitro* transcription template containing an optimized T7 promoter, 3’UTR, 5’UTR and a 100-adenine tail. The OVA, and IL-12 nucleoside modified mRNA sequences were prepared using the MegaScript transcription kit (ThermoFisher Scientific), co-transcriptionally capped using the 3’OMe CleanCap™ system (TriLink Biotechnologies) and purified using a modified cellulose base chromatography method (*34*), precipitated, eluted in nuclease free water, and quantified using the NanoDrop One system. Length and integrity were determined using the Agilent BioAnalyzer 2100 system. Endotoxin content was measured using the GenScript Toxisensor chromogenic assay, and values were below detection levels (0.1 EU/mL). mRNA was frozen at -20°C until formulation.

### Production and characterization of mRNA-LNP vaccines

Purified mRNAs were formulated into lipid nanoparticles using a self-assembly process wherein an ethanolic lipid mixture of an ionizable cationic lipid, phosphatidylcholine, cholesterol, and polyethylene glycol-lipid was rapidly combined with an aqueous solution containing mRNA at acidic pH as previously described (29). The ionizable cationic lipid (pKa in the range of 6.0-6.5, proprietary to Acuitas Therapeutics) and LNP composition are described in the patent application WO 2017/004143. The hydrodynamic size, polydispersity index (PDI) and zeta potential of mRNA-LNPs were measured using a Zetasizer Nano ZS90 (Malvern Instruments, Malvern, UK). The mRNA encapsulation efficiency was determined using a modified Quant-iT RiboGreen RNA assay (Invitrogen). Endotoxin levels were determined using the Limulus Amebocyte Lysate (LAL) chromogenic assay found to be <0.5 endotoxin unit (EU)/mL.

### Immunization

All mice were immunized intramuscularly (i.m.) in their hind legs with the indicated dose of mRNA-LNP in 50 µ.L of dPBS or 50 µ.L dPBS alone as a vehicle control. When vaccinating mice with multiple LNPs, particles were mixed such that an equivalent dose of LNP and antigen was given between groups. For example, 0.5 µ.g of LNP-OVA was mixed with 0.5 µ.g of eLNP, and 0.5 µ.g of LNP-OVA was mixed with 0.5 µ.g of LNP-IL12 when comparing the adjuvant affect of LNP-IL12. Unless otherwise indicated, particles were mixed together at a 1:1 ratio by particle mass. When immunizing mice that received OT-I T cell transfer, vaccination was performed the day following transfer.

### In Vivo CTL

An equal mix of congenically distinct target cells that were either pulsed with SIINFEKL peptide for 1 hour at 37°C or left unpulsed were transferred intravenously via tail-vein injection into mice that had been vaccinated with LNP-OVA or LNP-OVA+IL12 30 days prior. The unpulsed cells were stained with a high concentration of cell-trace violet (Invitrogen, C34557; 1:2000 dilution), while the unpulsed cells were stained at a low concentration (1:40,000 dilution) of cell trace violet to allow visual differentiation of the two populations. 16 hours post transfer the mice were sacrificed and target cell killing was assessed as the ratio of pulsed cells to unpulsed cells.

### T-cell transfers

For T cell transfers, OT-I transgenic mice were bred in in the University of Pennsylvania Department of Pathobiology vivarium in accordance with institutional guidelines. To isolate OT-I CD8^+^ T cells, we harvested spleens, and leukocytes were obtained by processing spleens over a 70-µm filter (Fisher Scientific, 22-363-548) and washing them in RPMI (Corning, 10-040-CM) with 5% FBS (RP-5). Red blood cells were then lysed by incubating for 3 min at room temperature in 5 mL of lysis buffer [0.864% ammonium chloride (Sigma-Aldrich, A0171) diluted in sterile deionized H_2_O], followed by washing with RP-5. For transfer of OT-I CD8^+^ T cells into wild-type hosts, splenocytes were rinsed with PBS, and total splenocytes containing 1 × 10^3^ OT-I T cells in 100 mL of dPBS were transferred retroorbitally. For the transfer of OT-I CD8^+^ T cells into non-WT mice, cells were enriched by magnetic activated cell sorting (MACS) using the CD8a^+^ T Cell Isolation Kit (Miltenyi Biotec, 130-104-075), and 1 × 10^3^ OT-I T cells in 100 mL of dPBS were transferred by retroorbital injection into recipient mice.

### Tissue preparation for Flow Cytometry

Leukocytes from the spleen and dLNs were obtained by processing spleens and LNs over a 70 µ m filter, washing them in complete media, and lysing red blood cells (see above). Cells were then resuspended in RP-5 (see above). Leukocytes from the lung were obtained by digesting lungs in RP-5 containing Type I collagenase (400 U/mL) and DNase (0.33 mg/mL) for 30-40 minutes at 37°C. Lungs were then processed over a 70 µm filter and washed with RP-5. Red blood cells were lysed (see above) and the cells were resuspended in RP-5.

### Flow Cytometry

Following processing, cells were stained for cell death using Ghost Dye Violet 510 Viability Dye (Tonbo Biosciences, 13-0870-T100), or Ghost Dye Red 780 Viability Dye (Tonbo Biosciences, 13-0865-T100) for 20 minutes at 4°C. Cells were washed with FACS buffer [1 × PBS, 0.2% bovine serum albumin (Gemini, 700-100P), and 1 mM EDTA (Gibco, 15575-038)] and incubated in Fc block [99.5% FACS buffer, 0.5% normal rat immunoglobulin G (Invitrogen, 10700), and 2.4G2 (1 µ g/ml; Bio X Cell, BE0307) and Mouse IgG isotype (3 µ g/mL; Thermofisher Scientific, 10400C)] at 4°C for 10 min before staining. Tetramer-specific CD8^+^ and CD4^+^ T cells were measured by staining in 50 µ L of FACS buffer containing MHCI tetramer for SIINFEKL for 35-45 minutes at 4°C. Cells were surface-stained in 50 µ L at 4°C for 20 to 25 min and washed in FACS buffer before acquisition. For detection of IL-12p40 protein in draining lymph nodes after vaccination with LNP-IL12, cells were replated in a 96-well plate in media containing Brefeldin A (5.0 mg/mL: Biolegend, 420601) and Monensin protein transport inhibitors (4 µ L/6E6 cells; GolgiStop, BD Biosciences, 554724) and incubated for 4 hours at 37°C. For intracellular cytokine staining of restimulated splenocytes, 4×10^6^ cells from spleen were plated in a 96-well plate and incubated with SIINFEKL peptide and anti-CD107a for 2 hours at 37°C before Brefeldin A (5.0 mg/mL: Biolegend, 420601) and Monensin protein transport inhibitors (4 pL/6E6 cells; GolgiStop, BD Biosciences, 554724) were added to the media. Cells were incubated for an additional 2 hours at 37°C before surface staining. For transcription factor staining, cells were rinsed with FACS buffer and surface-stained as described above, fixed using the eBioscience Foxp3 Transcription Factor Fixation/Permeabilization Concentrate and Diluent (Thermo Fisher Scientific, 00-8222) for 30 min at 4°C, and then washed with eBioscience Permeabilization Buffer (Thermo Fisher Scientific, 00 8333-56). Cells were then stained for transcription factors in 50 µ l of 1 × eBioscience Permeabilization Buffer at 4°C for at least 1 hour. Cells were then washed in eBioscience Foxp3 Transcription Factor permeabilization buffer and resuspended in FACS buffer before acquisition. For intracellular cytokine staining, cells were rinsed with FACS buffer and surface-stained as described above and fixed using the BD cytofix/cytoperm (BD Biosciences, 554714) permeabilization/fixation solution for 20 minutes at 4°C. Cells were then washed in BD perm/wash buffer and stained for intracellular cytokines in perm/wash buffer for 45-60 minutes at 4°C. Cells were then washed in FACS buffer before acquisition.

### Cell culture and BMDM assays

Bone marrow was harvested from the femur and tibia of WT or IL12p40^-/-^ mice and washed with complete DMEM (DMEM, 10% FBS, HEPES, Sodium Pyruvate, Pen/Strep). Red blood cells were lysed (see above) and cells were resuspended in complete DMEM containing 30% L929-supernatant (L-sup). Bone marrow cells were grown at 37°C, 5% CO2, and 7-9 days later, bone marrow derived macrophages (BMDMs) were used for LNP experiments or frozen down for future use at -80°C. BMDMs were frozen down at 1×10^7^ cells/mL in 90% FBS and 10% DMSO. Cells were frozen down to -80°C and 2-3 days later moved to -120°C. To thaw BMDMs, BMDMs were thawed at 37°C until just thawed and immediately washed with complete DMEM. Thawed cells were plated and allowed to recover in complete DMEM with 30% L-sup for 3 days at 37°C, 5% CO2 before LNP experiments were performed. For LNP experiments, 100 µ L of BMDMs at a concentration of 1.5×10^6^ cells/mL were plated in each well of a 96-well plate. 50 µ L of media containing the indicated dose of LNP was added to BMDM cultures and incubated overnight. To detach cells for flow cytometry staining, media was removed and wells were washed with 1X dPBS followed by addition of 0.25% Trypsin-EDTA (Gibco, 25200056) and incubation at 37°Cfor 5 minutes. Cells were gently triturated to remove them from wells and triplicate wells were pooled for flow cytometry staining (see above).

### ELISA

#### Draining lymph node lysates

Samples were kept on ice the entirety of the following process. The popliteal and inguinal draining lymph nodes were harvested and put into 600 uL of EZLys Tissue Protein Extraction Reagent (BioVision, 8002-500) containing 1X protease and phosphatase inhibitor cocktail (Thermofisher Scientific, 78440). Lymph nodes were mechanically lysed using a mini bead beater, set to ΨhomogenizeΨ for 90 seconds. After homogenizing, the samples were aliquoted and frozen down at -80°C until they were to be assayed. *BMDM supernatants*. Following overnight incubation (16-20 hours) with mRNA-LNPs, BMDMs and their supernatants were directly frozen down at 80^°^C until they were to be assayed. Draining lymph node and BMDM samples did not undergo more than one round of freeze-thaw. IFN-α production was assayed using the R&D biosystems mouse IFN-α all subtype Quantikine ELISA kit (MFNAS0) according to manufacturer’s instructions. To assay IL-12 production, a 4HBX Immulon 96-well plate (ThermoFisher, 3855) was coated with 0.5 µg/mL anti-IL12/23p40 clone 15.6 (Biolegend, 505201) in 1X dPBS at 4°C overnight. Standards were prepared by diluting recombinant IL12p70 (Peprotech, 210-12) in assay media at a max concentration of 20,000 pg/mL with 2-fold serial dilutions. Samples were added at the appropriate dilution in assay media and incubated for 2 hours at 37°C. 100 µ L of biotinylated anti-IL12/23p40 clone 17.8 (Biolegend, 505301) in 1x dPBS + 0.1% BSA was added at a concentration of 1 µ g/mL and incubated for 1 hour at RT. Horseradish peroxidase conjugated to avidin (HRP-Avidin; Biolegend, 405103) diluted 1000-fold in 1x dPBS + 0.1% BSA was added at 100 µ L/well. The plate was developed using 100 µ L of ABTS peroxidase substrate (SeraCare, 5120-0041) per well and development was stopped using an equal volume of 1% sodium dodecyl sulfate solution. The ELISA was read at 450 nm. Between each step, the plate was washed at least 5 times with 1X dPBS containing 0.05% Tween.

### Tumor Cell Lines

The OVA-secreting B16F0-OVA cell line was provided by Dr. Drew Weissman (University of Pennsylvania, Philadelphia, USA). Tumor cells were grown in RPMI-1640 containing 10% FBS, 10 mM HEPES, 1 mM sodium pyruvate, 1x GlutaMAX™, 1x MEM Non-Essential Amino Acid, 55 µ M β -mercaptoethanol, 1x Penicillin/Streptomycin G and G418 (400 µ g/ml) *(InvivoGen)*. ^56^

### Melanoma tumor model

The right flanks of study mice were shaved prior to tumor implantation to visualize tumor growth. 2 × 10^5^ B16F0-OVA cells were injected subcutaneously (s.c.) into the flank in 200 µ l of sterile Hanks’ Balanced Salt Solution (Gibco). For immunization, two doses of 10 µ g of indicated mRNA-LNPs were administered i.m. on day 4 and 12 after tumor inoculation. The growth of tumors was monitored every 2-3 days for at least 40 days or until humane endpoint was reached. Tumor volumes were monitored by using a vernier caliper and calculated using the equation: V = (4 × 3.14 × A × B^2^)/3, where V = volume (mm^3^), A = the largest diameter (mm), and B = the smallest diameter (mm). Mice were sacrificed when tumor size reached 4000 mm^2^ or if the tumors ulcerated.^56^

### UMAP analysis

The FlowJo UMAP plug-in version 4.0.3^76^ was used to generate UMAP projections of flow cytometry data. Prior to UMAP analysis, keywords were used to label features of interest for each sample (ex: tissue, timepoint, treatment) in order to preserve them during concatenation, and populations of interest were concatenated using FlowJo’s concatenate function. UMAP analysis was then run using all parameters except those used for negative selection.

### Statistics

Statistics were performed using GraphPad Prism Version 10.2.2 statistical analysis software. Normality and log-normality tests were carried out prior to testing. For normally distributed data, comparing two groups along a single variable, a standard Student’s T-test was used. If the data was found to be abnormally distributed, a Mann-Whitney Rank Sum Test was used. In the comparison of two groups along more than one variable, a Two-Way ANOVA with a follow-up Šídák’s multiple comparison test was used. P-values less than 0.05 were considered significant.

---

**Supplementary Figure 1.**
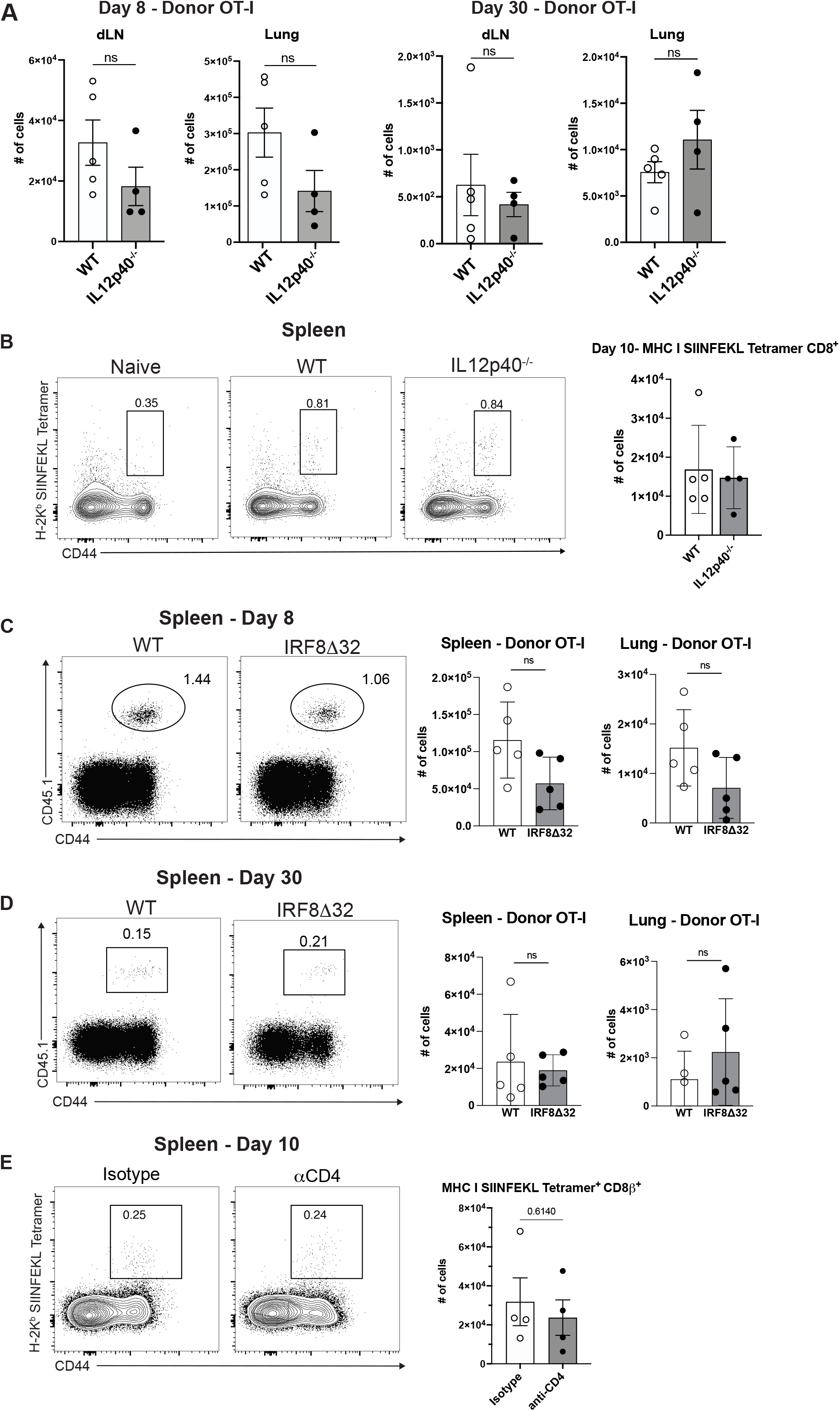
IL12p40^-/-^ and related pathways are not required for CD8^+^ T cell responses to mRNA vaccination. **(A)** Flow cytometry analysis of OT-I T cell numbers in draining LNs and lungs of mice at day 8 (left) and 30 (right) post immunization with LNP-OVA. Summary plots show OT-I T cell number in WT (open circles, open bars) and IL12p40^-/-^ (closed black circles, gray bars) mice in indicated tissues and at indicated time points. **(B)** Flow cytometry analysis of the endogenous CD8^+^T cell response to LNP-QVA vaccination 10 days post immunization. Representative flow cytometry plots (left) and summary plots (right) show the total number of splenic CD8^+^ T cells that stained positive for the H2-K^b^:SIINFEKL Tetramer in WT (open circles, open bars) and IL12p40^-/-^ (closed black circles, gray bars) mice. **(C-D)** Flow cytometry analysis of OT-I T cell numbers in the spleen (left) and lung (right) of mice at day 8 (C) and 30 (D) post immunization with LNP-OVA. Summary plots show OT-I T cell numbers in WT (open circles, open bars) and cDC1 deficient, IRF8A32 (closed black circles, gray bars) mice in indicated tissues. **(E)** Flow cytometry analysis of the endogenous CD8^+^T cell response to LNP-OVA vaccination 10 days post immunization. Representative flow cytometry plots (left) and summary plots (right) show the total number of splenic CD8^+^ T cells that stained positive for the H2-K^b^:SIINFEKL Tetramer in isotype treated (open circles, open bars) and CD4 depleted (closed black circles, gray bars) mice. All summary plots show mean and standard error of the mean. A Shapiro-Wilk normality test was carried out prior to testing for (A-E) followed by an unpaired student’s T-Test. Data is shown from 1 replicate of an experiment performed 3 times for (A), 1 replicate of an experiment performed 1 time for (B-E).

**Supplementary Figure 2.**
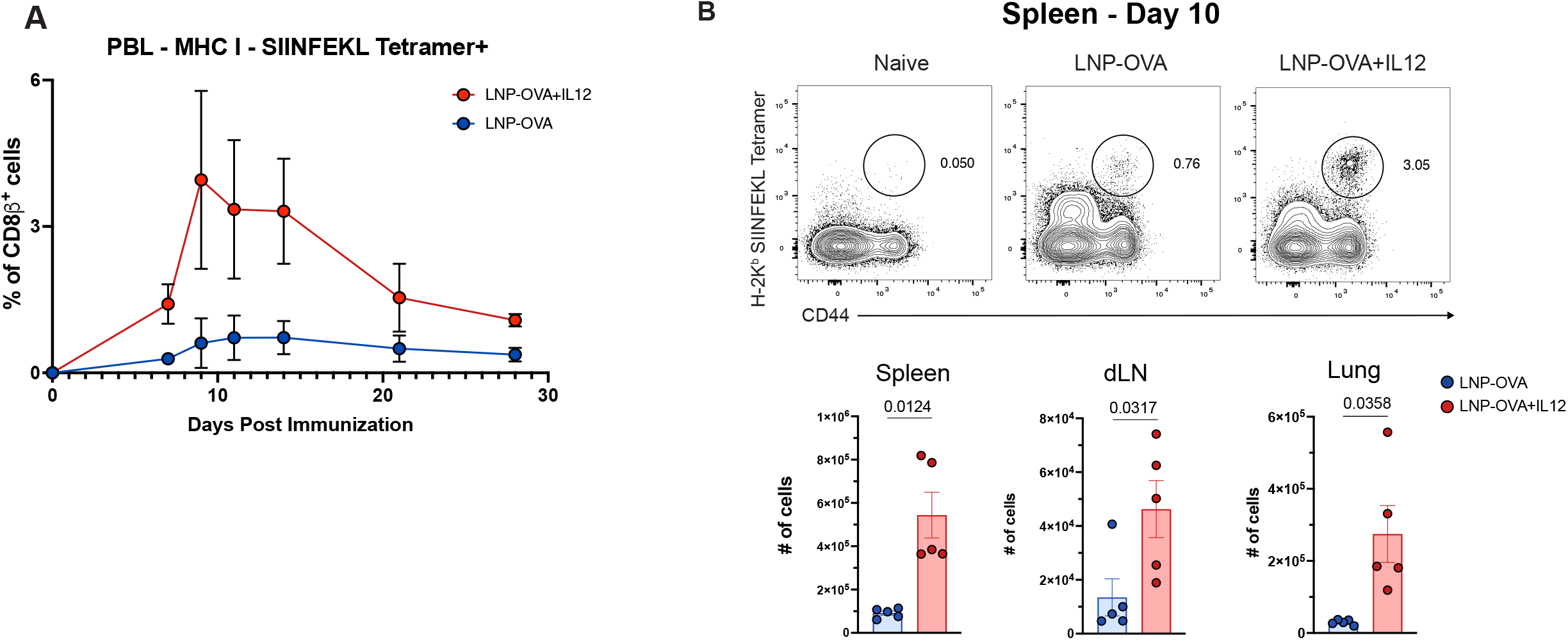
**(A-B)** Flow cytometry analysis of the endogenous CD8^+^T cell response to LNP-OVA and LNP-OVA+IL12 vaccination. **(A)** Frequency of H2-K^b^:SIINFEKL CD8^+^ T cells in blood at days 7, 9, 11, 14, 21 and 28 post immunization with LNP-OVA (blue) LNP-OVA+IL12 (red). **(B)** Flow cytometry analysis of the endogenous CD8^+^T cell response to LNP-OVA vaccination 10 days post immunization. Representative flow cytometry plots show the total number of splenic CD8^+^ T cells in the spleen (left), dLN (center), and lung (right) that stained positive for the H2-K^b^:SIINFEKL Tetramer in LNP-OVA (blue) and LNP-OVA+IL12 (red) mice. Summary plots show mean and standard error of the mean. A Shapiro-Wilk normality test was carried out prior to testing for (B) followed by a Mann-Whitney rank sum test. Data shown for one replicate of an experiment performed 3 times.

**Supplementary Figure 3.**
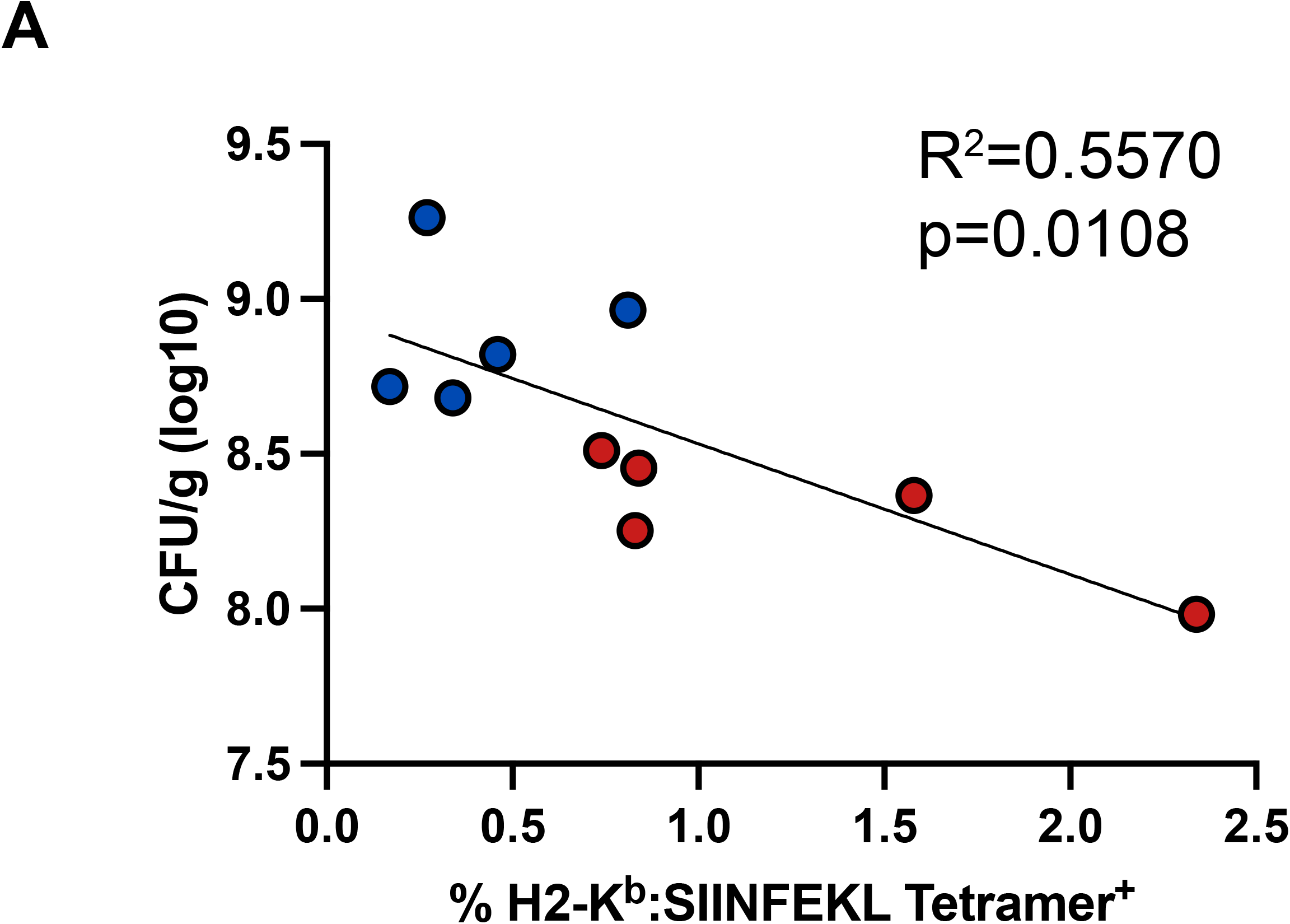
**(A)** Correlation between pre-challenge OVA-specific CD8^+^ T cell frequency in the blood and post challenge bacterial colonization in the spleen. Mice were vaccinated with LNP-OVA (blue) or LNP-OVA+IL12 (red) and bled 26 days post vaccination, then challenged 1 week later. Simple linear regression was used to determine R^2^ and significance values. Data shown for 1 replicate of an experiment performed 2 times.

